# Uncertainty Modeling Outperforms Machine Learning for Microbiome Data Analysis

**DOI:** 10.1101/2025.09.12.675956

**Authors:** Maxwell A. Konnaris, Manan Saxena, Nicole Lazar, Justin D. Silverman

## Abstract

Microbiome sequencing measures relative rather than absolute abundances, providing no direct information about total microbial load. Normalization methods attempt to compensate, but rely on strong, often untestable assumptions that can bias inference. Experimental measurements of load (e.g., qPCR, flow cytometry) offer a solution, but remain costly and uncommon. A recent high-profile study proposed that machine learning could bypass this limitation by predicting microbial load from sequencing data alone. To evaluate this claim, we assembled *mutt*, the largest public database of paired sequencing and load measurements, spanning 35 studies and over 15,000 samples. Using *mutt*, we show that published machine learning models fail to generalize: on average they perform worse than a naive baseline that always predicted the training set mean. These failures stem from covariate shift–limited shared taxa between studies, differences in community composition, and differences in preprocessing pipelines–that silently derail model inputs. In contrast, Bayesian partially identified models do not attempt to impute microbial load, but instead propagate scale uncertainty through downstream analyses. Across 30 benchmark datasets, Bayesian partially identified models consistently outperformed normalization and machine learning approaches, providing a principled and reproducible foundation for microbiome inference.

## INTRODUCTION

Microbiome sequencing enables high-throughput, culture-independent profiling of microbial communities^1–3^. Yet sequencing measures only relative, not absolute, abundances: sequencing depth is unrelated to total microbial load^4–6^. This limitation undermines many quantitative analyses^6–9^. To address this, sequence counts are often normalized as part of statistical analyses^10^. However, recent studies have shown that these normalization methods imply strong, often unverifiable, assumptions about microbial load^9^. For example, total-sum scaling (TSS) implicitly assumes that all communities contain the same total number of microbes^7,8^. Evaluations using datasets with paired sequencing and microbial load measurements (e.g., qPCR or flow cytometry) have revealed the consequences of these assumptions: even small violations can inflate false discovery rates beyond 75% and reduce statistical power^7,8,11^.

Two alternatives have recently been proposed. Nixon et al. ^9^ reframed the problem of unknown total microbial load (i.e., “unknown scale”) as one of partially identified statistical models. They introduced a class of Bayesian partially identified models (PIMs) that explicitly represent scale uncertainty, rather than imposing artificial identifiability through normalization. Follow-up studies have shown that modeling this uncertainty directly can improve statistical power while dramatically reducing false discovery rates, even under substantial violations of scale-related assumptions^7,8,11–14^.

More recently, Nishijima et al. ^15^ proposed an alternative strategy: using machine learnin (ML) to predict total microbial load directly from sequencing data. Their models were trained on four previously published datasets with paired sequencing and flow cytometry measurements and reportedly achieved a correlation of approximately 0.6 between predicted and measured microbial load samples on held-out data. From this, the authors claim that their models generalize to new datasets. The authors applied their models to more than 150 additional studies which lacked microbial load measurements, and reported ML-based predictions uncovered novel biological patterns missed by existing methods. These models have since gained substantial attention and are being adopted in ongoing studies^16^.

In this article, we critically evaluate the generalizability of these ML-based microbial load prediction models. We show that the sample correlation metric used by the original authors is misleading: more appropriate measures reveal that even on the three original datasets performance is weak (coefficient of determination, *R*^2^, between predicted and measured loads of ≈ 0.3, meaning the models explain only about 30% of the variance). Moreover, slight changes in preprocessing or preservation cause accuracy to collapse, with *R*^2^ < 0–worse than a naive model that simply predicts the training set mean. To move beyond these limited evaluations, we assembled the *mutt* database, comprising 35 studies and over 15,000 samples with paired sequencing and microbial load. Using this resource, we find that the published ML models consistently fail on independent datasets across diverse populations, protocols, and pipelines, again performing worse than mean prediction and even producing predictions that were anti-correlated with measured microbial loads. We show that these failures stem from covariate shift–differences between the distribution of features in the training set and the test sets. We identify multiple factors driving the covariate shift including limited taxonomic overlap across studies: models often expect taxa that are absent in new datasets while ignoring novel taxa, forcing inputs to be zero-filled and causing predictions to collapse. Our analyses of the *mutt* database also provide the most comprehensive assessment of normalization-based methods to date, confirming false discovery rates often exceed 75% and positive predictive values fall below 25%. Finally, we show that Bayesian PIMs, which model scale uncertainty rather than predict microbial load, consistently outperform both ML- and normalization-based methods in differential abundance tasks, offering a more robust foundation for microbiome analysis.

## RESULTS

### Reproducing Model Performance on Held-out Datasets

We re-evaluated the predictive performance of the ML models developed by Nishijima et al. ^15^. Their study trained separate models on three gut microbiome cohorts with paired microbial load and sequencing data: two shotgun metagenomic datasets (MetaCardis, *n* = 1,812^17^ and GALAXY, *n* = 1,894^15^) and one 16S rRNA dataset (Vandeputte 2021, *n* = 707^18^). The MetaC-ardis and GALAXY models were cross-validated on each other, while the Vandeputte (2021) model was tested on an earlier 16S rRNA cohort from the same group (Vandeputte 2017; the *disease cohort*). Nishijima et al. ^15^ report correlations of *r* ≈ 0.56–0.60 between predicted and measured microbial load in these validation studies.

Using the relative abundance tables provided by the authors, we reproduced their reported correlations: *r* = 0.56 for GALAXY, *r* = 0.56 for MetaCardis, and *r* = 0.60 for the Vandeputte (2021) model on their respective held-out datasets. Moreover, reprocessing raw reads from the Vandeputte (2017) study per the authors stated methods, the only held-out dataset with publicly available raw data, again gave the same result (sample correlation of *r* ≈ 0.61).

While reproducible, we found the authors’ reliance on sample correlations to evaluate performance inadequate. Correlation only measures whether predictions increase when the true values increase, and can still be high even when the model predicts nearly the same value for every sample, despite true microbial loads spanning orders of magnitude (Supplemental Figure 1). Such compressed predictions can bias downstream analyses like differential abundance, which depend on the size of between-sample differences. To better assess predictive utility, we instead computed the mean-centered coefficient of determination (*R*^2^), which quantifies the proportion of variance explained and penalizes both biased and low-variance predictions. Because the Nishijima et al. ^15^ models were trained on flow cytometry-based measurements, but often validated on datasets measured with different modalities (e.g., qPCR, ddPCR), we mean-centered predictions before calculating *R*^2^ so that systematic offsets between measurement types would not artificially reduce performance. Note that negative *R*^2^ values indicate that a model performs worse than a naive baseline that simply predicts the mean microbial load across all samples.

Quantifying model performance using the mean-centered coefficient of determination showed that the Nishijima et al. ^15^ models had limited predictive performance: *R*^2^ of 0.28, 0.30, and 0.31 for the MetaCardis, GALAXY, and Vandeputte (2021) models on their respective held-out datasets (Figure 1A; Supplemental Table 2). Thus, although we reproduced the correlations reported by Nishijima et al. ^15^, re-evaluation with a more appropriate metric revealed that their models explain less than one-third of the variance in microbial load.

**Figure 1:**
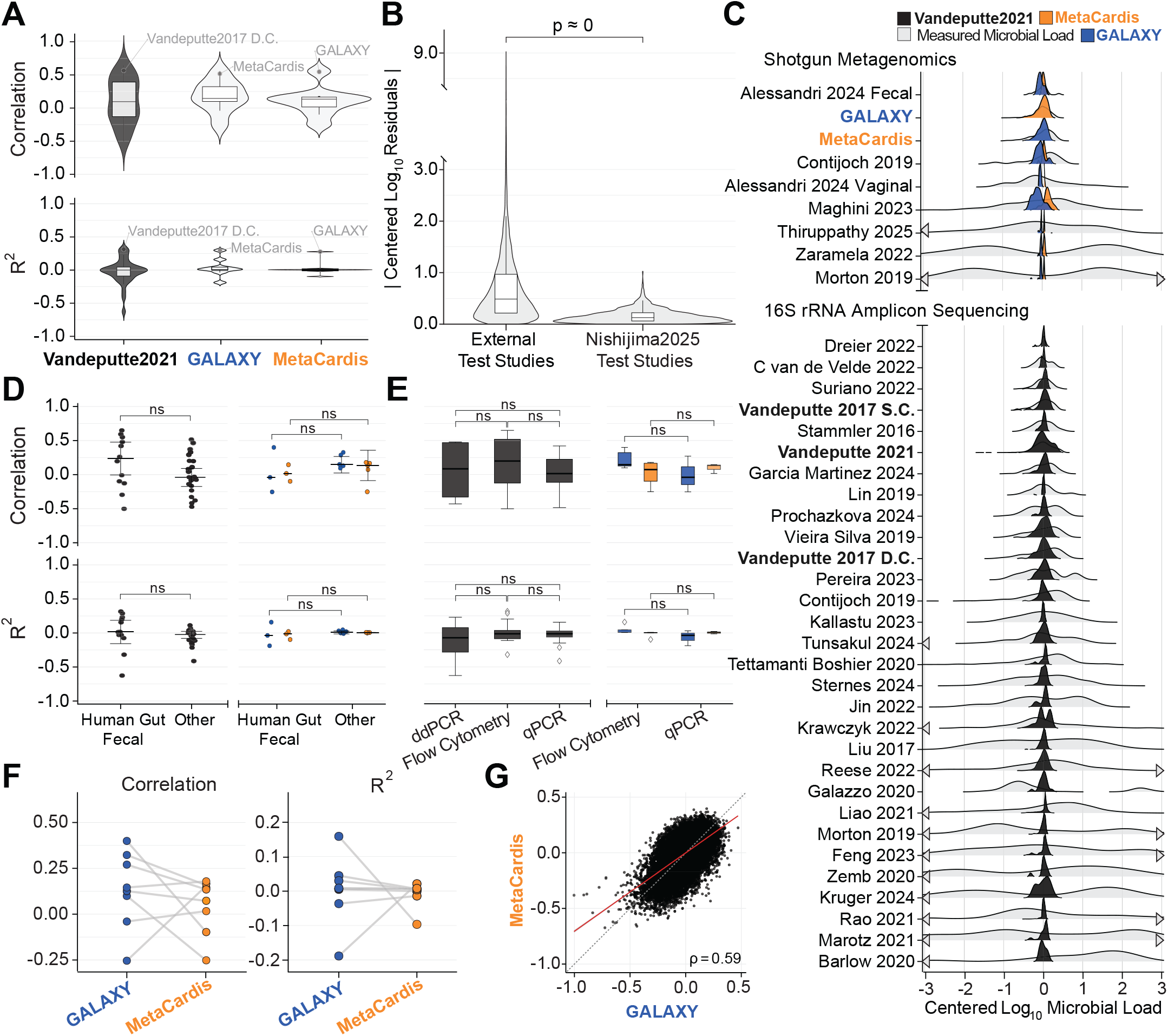
Nishijima et al. ^15^ microbial load prediction models fail to generalize out of distribution. **(A)** Performance of each of the three Nishijima et al. ^15^ models (x-axis) applied to the three original test datasets from that study (labeled points) and 31 external datasets from the *mutt* database. Sample correlation (*r*) and meancentered coefficient of determination (*R*^2^) were calculated within each study for 16S rRNA (dark gray) and shotgun metagenomic (light gray) sequencing data. By definition, a baseline predictor that always predicts the mean of the training set will have *R*^2^ = 0, this is indicated by a dashed line. **(B)** Distribution of absolute mean-centered log_10_ residuals for samples from the *mutt* database (labeled “External Test Studies”) compared to the original test datasets (labeled “Nishijima2025 Test Studies”). **(C)** Measured microbial loads (gray) and predictions from the GALAXY (blue), MetaCardis (orange), and Vandeputte2021 (black) models. Distributions are truncated between −3 and 3 log_10_ units; arrows indicate tails extend beyond plotted range. **(D)** Performance stratified by whether a samples was from human feces versus other biospecimen type (e.g., human saliva) for external datasets. **(E)** Performance stratified by microbial load measurement modality (flow cytometry, qPCR, ddPCR) for external datasets. **(F)** Study-level differences in *R*^2^ between predictions from the GALAXY and MetaCardis models. **(G)** Sample-level concordance between mean-centered log_10_ predictions from the GALAXY and MetaCardis models applied to the “global datasets” analyzed in Nishijima et al. ^15^ (*N* = 164; *n* = 27,832 samples). Red line shows ordinary least squares fit; dashed black line represents 1:1 agreement. Sample correlation *r* is shown in the lower-right. Boxplots indicate median ±IQR with whiskers at 1.5×IQR. All comparisons were performed using Wilcoxon rank-sum tests (ns: not significant). *N*, number of studies.

Going beyond the original Nishijima et al. ^15^ analysis, we hypothesized that technical factors affecting model features, such as differences in sample preservation or data preprocessing, would impair predictive performance. To test this hypothesis we analyzed an additional *study cohort* (*N* = 45) from Vandeputte et al. ^6^ which was not analyzed in Nishijima et al. ^15^. This *study cohort* includes flow cytometry measurements from both frozen and fresh samples whereas the *disease cohort* analyzed in Nishijima et al. ^15^ only had measurements from frozen samples. On this additional cohort, the Vandeputte (2021) model performance was low and preservation-dependent: *R*^2^ ≈ 0.09 on frozen specimens versus *R*^2^ ≈ 0.00 on fresh specimens, with all other preprocessing held constant.

We also repeated the analysis using the count tables from the original Vandeputte et al. ^6^ study, rather than the reprocessed tables provided by Nishijima et al. ^15^, reclassifying taxa with the RDP Classifier v16 to mirror the methods of Nishijima et al. ^15^. Using this alternative preprocessing pipeline, performance worsened in the disease cohort (frozen; *R*^2^ ≈ 0.12) and became negative in the study cohort (frozen: *R*^2^ ≈ − 0.11; fresh: *R*^2^ ≈ −0.14; Supplemental Table 2).

Taken together, these analyses demonstrate that the published findings of Nishijima et al. ^15^ are reproducible from their released data and models. However, even within these carefully matched test sets, predictive performance remained limited (*R*^2^ ≈ 0.3), and modest changes in sample preservation, preprocessing, or taxonomic classification pipelines further eroded accuracy. These observations highlight the fragility of the Nishijima et al. ^15^ microbial load prediction models and raise fundamental concerns about their robustness and generalizability.

### *mutt* : A Curated Public Database of Microbiome Studies with Paired Sequencing and Microbial Load Measurements

Motivated by the limited scope of the original evaluation in Nishijima et al. ^15^, where each model was tested on only a single held-out dataset, we assembled *mutt*: a curated, publicly available database of studies that pair sequence count data with direct measurements of microbial load. *mutt* is designed to advance rigor and reproducibility in microbiome research.

The *mutt* database spans 35 studies (>15,000 samples) covering diverse sample types (e.g., human oral, fecal, and semen; multi-species gut; soil; wastewater; cheese) and sequencing modalities (shotgun metagenomics and amplicon sequencing). Each study is paired with external microbial load measurements (e.g., flow cytometry, qPCR, ddPCR, CFU counts, spike-in controls) and associated metadata. To our knowledge, mutt is the most comprehensive curated resource for studying absolute microbial abundances to date.

For studies with available raw data, *mutt* supplements published count tables with uniformly processed versions generated using MetaPhlAn4^19^ (metagenomics) or DADA2^20^ (amplicons). Supplemental Table 1 details the 125 studies screened, outcomes of data requests where files were missing or embargoed, and the final subset included in this analysis.

All code, processed data, and issue tracking for *mutt* are publicly available at https://github.com/Silverman-Lab/mutt.

### Predictive Performance Collapses Outside of Training Domain

In addition to the three held-out test sets evaluated by Nishijima et al. ^15^, we applied their published models to 31 external studies from the *mutt* repository that met our inclusion criteria (see Methods; Figure1A; Supplemental Table1). Together, these studies contributed 34 datasets: seven shotgun metagenomic datasets were used to test the GALAXY and MetaCardis models, and 27 16S rRNA amplicon datasets were used to test the Vandeputte (2021) model. The total includes two studies that provided both shotgun and amplicon data, as well as one shotgun study that yielded separate fecal and vaginal datasets.

Across all external studies, predictive performance was poor. In over half of the datasets, the Nishijima et al. ^15^ models performed worse than a baseline predictor which always predicted the mean of the training set (*R*^2^ < 0). The overall mean ± standard deviation was 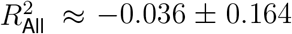 and sample correlation *r*_All_ ≈ 0.09 ± 0.32. Model-specific results were similarly weak: 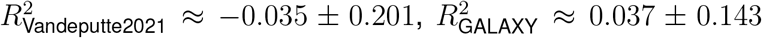, and 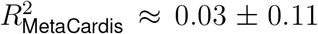 (Figure 1A; Supplemental Table 2). For many studies, the Nishijima et al. ^15^ models were anticorrelated with measured loads: the model predicted high loads for samples with low loads and vice versa. Performance across all external studies was significantly worse than on the three held-out datasets used in Nishijima et al. ^15^ (*p* ≈ 0; two-sided Wilcoxon-rank sum test, applied to absolute mean-centered log_10_ residuals; Figure 1B). Moreover, model predictions on external datasets were compressed: they had a much narrower range than measured microbial loads (Figure 1C). In many cases, the Nishijima et al. ^15^ models predicted nearly identical loads for almost every sample in a study (Supplemental Figure 2). Together, these results demonstrate that the Nishijima et al. ^15^ models do not generalize to new datasets.

We hypothesized that the Nishijima et al. ^15^ models might perform better on the 13 external human gut microbiome datasets since these matched the biological site of the training data. This subset included three shotgun metagenomic and ten 16S rRNA amplicon sequencing datasets. Even within this matched setting, model performance remained poor. The overall mean ± standard deviation was 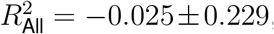, with model-specific values of 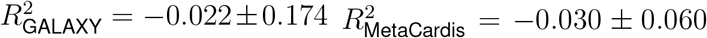, and 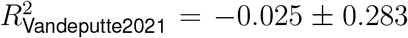 (Figure 1D; Supplemental Table 2). Moreover, predictive performance did not differ significantly by measurement modality: results were similarly poor across studies using flow cytometry, ddPCR, or qPCR for microbial load estimation (Figure 1E; Supplemental Table 2). Full results, including *R*^2^ and sample correlation *r* for each model across all studies, are provided in Supplemental Table 2. Together, these findings indicate that the models failed not only in out-of-domain contexts, but also within the same sample types, measurement platforms, and sequencing modalities on which they were originally trained.

Finally, performance of the two shotgun metagenomics models (GALAXY and MetaCardis) was inconsistent across datasets. In some instances, the GALAXY model’s *R*^2^ was ≈ 0.26 higher than the MetaCardis’, but in others, it was ≈ 0.21 lower. Thus there is no clear criterion for model selection (Figure 1F; Supplemental Table 2). We further assessed per-sample model agreement between GALAXY and MetaCardis trained models using the 164 global datasets provided by Nishijima et al. ^15^ (Figure 1G). These datasets do not have measured microbial loads so we are limited to assessing the concordance of the two models predictions. Across the 27,832 samples in the global dataset, the GALAXY and MetaCardis models had only moderate correlation (sample correlation *r* = 0.59 between mean-centered log_10_ predicted microbial loads). Taken together, the large variability in *R*^2^ and only moderate agreement between models indicate that there is no reliable basis for choosing which model to apply, further underscoring their lack of generalizability and practical utility.

### Covariate Shift Drives Poor Generalization of Nishijima *et al*. Models to External Data

We hypothesized that the observed decline in predictive accuracy arises from covariate shift. Covariate shift occurs when the distribution of input features–here, microbial relative abundances– differs between the training and test data of a machine learning model. Even if the underlying biological relationship between features and target remains constant, such distributional mismatches can substantially degrade performance on new populations or conditions^21^. This problem is acute in microbiome studies, where community composition varies widely across host populations, geographic regions, diets, and sequencing protocols, and where few taxa are consistently shared across studies^22^. As a result, models trained on taxonomic counts from one population are almost guaranteed to face covariate shift in another, even when sample type (e.g., stool) or measurement modality (e.g., flow cytometry) are held constant. In practice, we expect many taxa present in the training data are absent from the test data, forcing models to rely on incomplete or zero-filled feature vectors, which can cause systematic prediction failures.

Consistent with our hypothesis, a t-SNE projection of 16S rRNA samples from 27 *mutt* studies showed that model performance depended on a sample’s proximity to the training data. Samples from studies with high *R*^2^ values clustered near the training cohort in the embedding, while those from poorly performing studies lay further away (Figure 2A). One possible explanation is that studies conducted by the same or closely related research groups often involve comparable populations, protocols, and environments, leading to more similar microbial compositions. Supporting this, we found that external studies with overlapping first or senior authorship with the training cohort clustered more closely with the training data in feature space. Moreover, these studies exhibited significantly higher predictive performance than those without shared authorship (two-sided Wilcoxon rank-sum test, *p* = 0.0023, *N*_author-present_ = 6, *N*_author-absent_ = 23; Figure 2B). This comparison was restricted to amplicon data due to limited metagenomic data (*N*_author-present_ = 1, *N*_author-absent_ = 6; Figure 2B).

**Figure 2:**
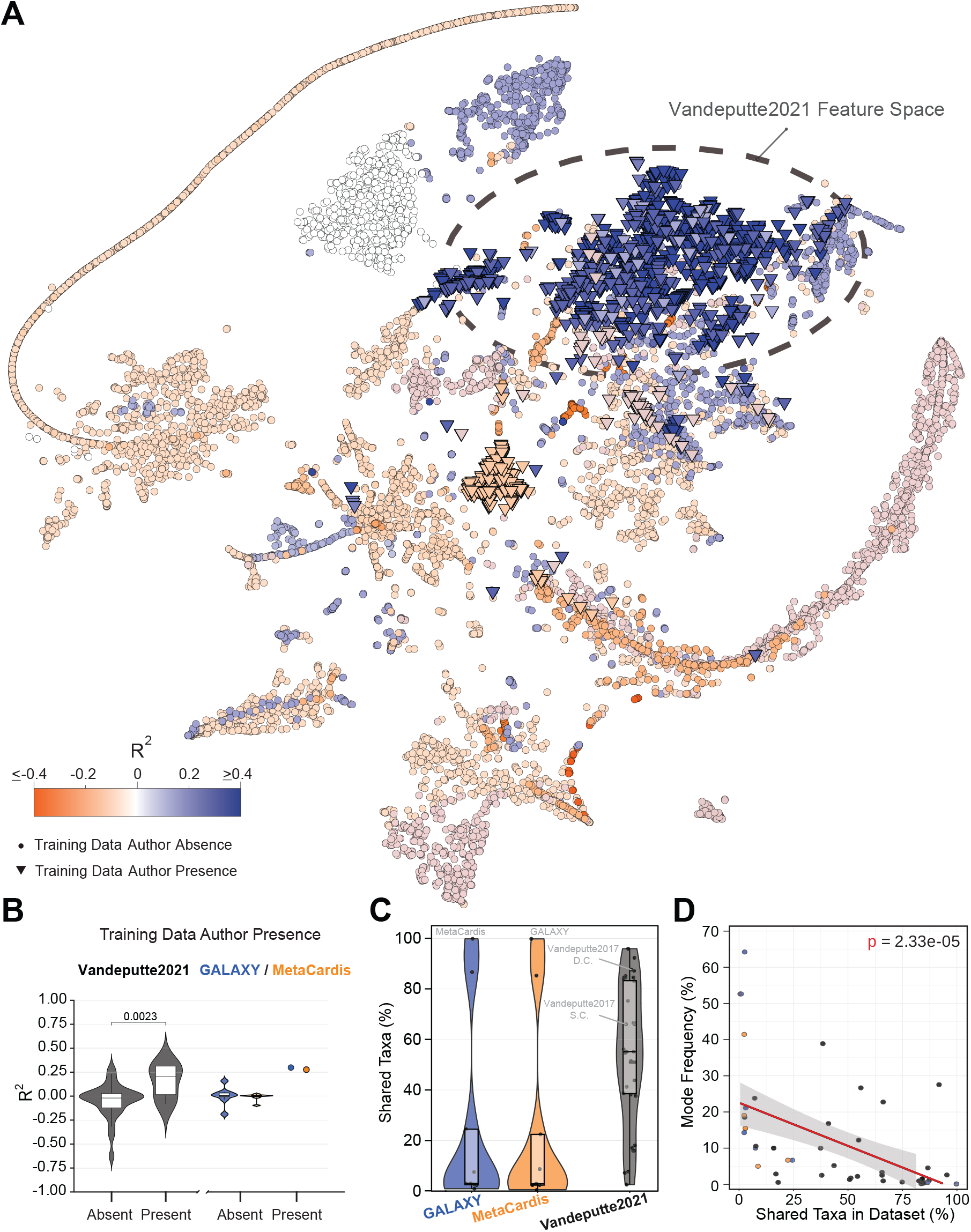
Covariate shift limits machine learning performance. **(A)** t-SNE projection of the 194 relativeabundance features used by the Vandeputte (2021) model ^15^ for amplicon datasets within the *mutt* database (*N* = 29). Points represent samples (*n* = 10, 933), shown as inverted triangles if the study shares a first or corresponding author with the training cohort, or circles otherwise. The dashed black ellipse marks the 99% confidence region of Vandeputte (2021) training samples ^18^ in feature space. Colors indicate study-level *R*^2^. **(B)** Study-level *R*^2^ by presence or absence of a first or corresponding author from any training study. Black, blue, and orange denote the Vandeputte (2021) amplicon, GALAXY shotgun, and MetaCardis shotgun models, respectively. Significance for the amplicon model was assessed via Wilcoxon rank-sum test (Author-present, *N* = 6; Author-absent, *N* = 23). Boxplots show median and IQR; whiskers extend to 1.5×IQR. **(C)** Percentage of taxa expected by a model that are present in a validation study. **(D)** Relationship between the frequency of the most common predicted value in a study (Mode Frequency) and percent of shared taxa per study. Red line depicts fitted linear model, with estimated slope of −0.23 and corresponding standard error of 0.05, corresponding p-value is shown as an annotation.

These findings suggest that covariate shift between the original training and independent datasets, not stochastic noise or simple overfitting, is the primary driver of model failure. Further inspection of the model architecture and input handling supports this interpretation. Each model expects a fixed-length input vector with specific taxonomic features (e.g., 194 taxa for the Vandeputte (2021) model). However, none of the 31 external datasets contained the full set of required taxa (Figure 2C). When expected taxa were missing, the model silently treated them as zeros, while taxa present in the external dataset but absent from the training set were ignored. In some cases (e.g.,van de Velde et al. ^23^, Dreier et al. ^24^, Alessandri et al. ^25^, Zaramela et al. ^26^, Morton et al. ^27^, Lin et al. ^28^, Thiruppathy et al. ^29^), fewer than 20% of the required taxa were available, meaning over 80% of the input covariates were treated as zeros. This extensive zero imputation likely drove the prediction collapse observed in Figure 1C and Supplemental Figure 2, where many datasets showed large fractions of samples assigned identical microbial load estimates. Supporting this, we observed a strong correlation between the proportion of shared taxa in a dataset and the frequency of the most common predicted value (Figure 2D).

We also observed that the original model implementations lacked input validation or clear format specifications, which allowed common preprocessing mismatches (e.g., taxonomic label misalignment) to silently produce erroneous inputs. Even after manual correction and alignment of taxonomic features, prediction collapse and poor generalization persisted—further underscoring that covariate shift, not user error or implementation flaws, lies at the heart of model failure.

Together, these findings suggest that the poor generalization of the Nishijima et al. ^15^ models is not due to model complexity or overfitting, but rather to an under-recognized covariate shift problem—one that is likely common when ML models are trained on taxonomic features.

### Bayesian PIMs Offer a Principled and Robust Alternative to Normalization and Scale Prediction

To evaluate how microbial load estimation influences downstream biological inference, such as differential abundance testing, we benchmarked different modeling strategies using 30 datasets from the *mutt* database with metadata-defined groups suitable for comparison (24 amplicon, six shotgun; see Supplemental Table 1).

We first evaluated widely used normalization-based methods, including DESeq2^30^, Limma-Voom^31^, ANCOM-BC2^32^, edgeR^33^, metagenomeSeq^34^, and LinDA^35^. For comparison, we turned to ALDEx3, an efficient extension of the ALDEx2 framework that provides a unified platform for differential abundance testing with customizable *scale models*. This allowed us to directly assess the impact of different approaches to modeling microbial load (see *Methods*). Specifically, we applied ALDEx3 with three alternative scale models: (i) Total Sum Scaling (TSS), which assumes no difference in microbial load between conditions^8^ (labeled “TSS” in Figure 3); (ii) the ML models from Nishijima et al. ^15^ (labeled “GALAXY”, “MetaCardis”, and “Vandeputte2021”); and (iii) a Bayesian prior model that generalizes TSS by incorporating uncertainty into the assumption of equal load (labeled “Bayesian PIM”). The Bayesian PIM corresponds to assuming, with 95% probability, that microbial load in one condition lies within a factor of 0.26–3.89 of the other, thereby explicitly propagating scale uncertainty into downstream inference.

**Figure 3:**
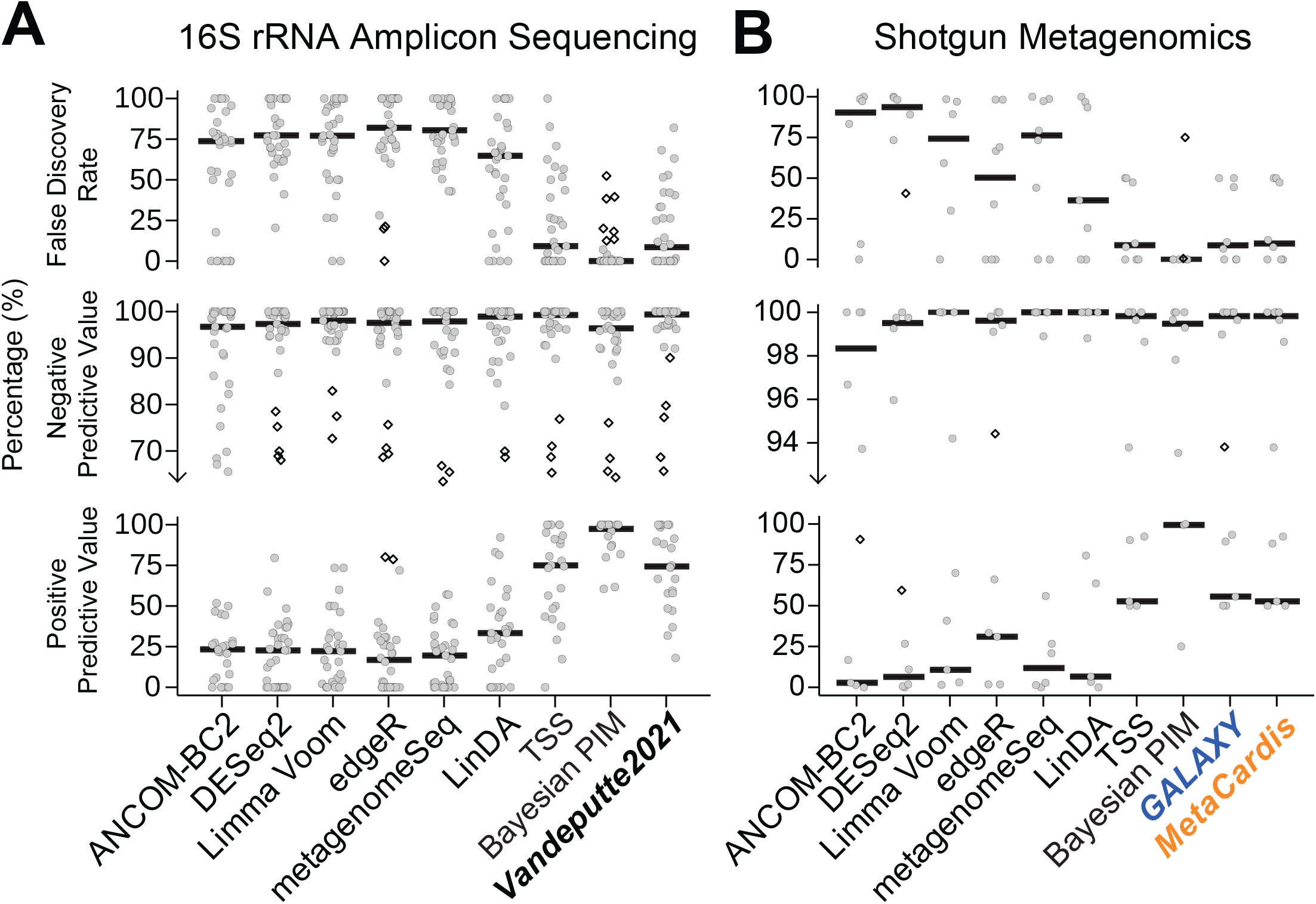
Bayesian PIMs outperform machine learning and normalization-based methods in downstream inference. **(A)** 16S rRNA amplicon (*N* = 24) and **(B)** shotgun metagenomic (*N* = 6) studies were used to evaluate multiple differential abundance methods. Multiple measurement modalities of microbial load (e.g. qPCR, flow cytometry, ddPCR) were included in the analysis per study. Approaches included ANCOM-BC2, DESeq2, Limma Voom, edgeR, metagenomeSeq, LinDA, and ALDEx3 with three different scaling strategies: (1) ALDEx3 with total sum scaling normalization (TSS), (2) ALDEx3 with relative abundances scaled by predicted microbial loads (Vandeputte2021, GALAXY, MetaCardis), and (3) ALDEx3 with a Bayesian PIM prior (i.e., scale model) that generalizes total sum scaling by considering uncertainty in scale (see *Methods*). Performance was assessed using three metrics: False Discovery Rate, Negative Predictive Value, and Positive Predictive Value with gray jittered points indicating study-level results and a solid black line representing the median of the distribution. Observations beyond 1.5 times inter-quartile range are plotted as diamonds. Ground truth was assessed using an ALDEx3 model scaled by measured microbial loads with a small amount of uncertainty introduced to account for measurement error (see *Methods*). Fold-change direction and Benjamini–Hochberg adjusted *p*-values were compared between each method and the ALDEx3 reference model to derive true positive, false positive, true negative, and false negative rates.

This final strategy exemplifies the Bayesian PIM framework proposed in Nixon et al. ^7^, reflecting a weakly informative prior grounded in biological plausibility—for example, that the human gut has limited carrying capacity and coarse homeostatic constraints. To our knowledge, with 30 datasets, this represents the most comprehensive real-data evaluation of scaling strategies for microbiome inference to date.

As expected from prior studies, normalization-based methods consistently failed to control the false discovery rate (FDR), yielding median type I error rates ≥ 75% and positive predictive values (PPV) < 25% (Supplemental Table 3). Using microbial load predictions from Nishijima et al. ^15^ as fixed inputs did not improve performance over simple TSS normalization: median FDR remained ≈ 9% and PPV ≈ 74% in amplicon datasets, nearly identical to the TSS model (Figure 3A). This is consistent with our earlier results (Figure 1C and Supplemental Figure 2), which showed that the Nishijima models often predict identical microbial loads with low variance across samples within a study. Unsurprisingly, then, they behave like TSS normalization which also assumes no systematic difference in microbial load between groups.

In contrast, the Bayesian PIM, which generalizes TSS by acknowledging uncertainty in how microbial load changes between conditions, delivered large and consistent improvements. The Bayesian PIM achieved near-perfect FDR control (median FDR≈ 0) while preserving statistical power (median PPV ≈ 99% and NPV ≈ 97%). These gains held across both amplicon and metagenomic datasets (Figure 3; Supplemental Table 3). Crucially, these improvements were not the result of model tuning or external calibration, but emerged from a more principled statistical treatment: explicitly acknowledging that total microbial load is unobserved leading to partially identified statistical models. Rather than substituting a single “best guess” for this unknown, Bayesian PIMs propagate uncertainty through the full inferential pipeline–enhancing reproducibility, reducing false positives, and maintaining power. This approach generalizes beyond microbiome studies to any setting where scale is unmeasured or uncertain, and it offers a more robust foundation for statistical inference than point-estimation-based methods^7,11^.

These results demonstrate that modeling uncertainty in scale can dramatically improve statistical performance in microbiome research.

## DISCUSSION

Machine learning promises to uncover biological insights from complex data—but only when its predictions generalize beyond the training context. Our analysis shows that microbial load prediction models trained on taxonomic counts, such as those from Nishijima et al. ^15^, fail to generalize to new populations, protocols, or processing pipelines. Even on held-out data from the same laboratories, these models explained less than a third of the variation in microbial load (*R*^2^ ≈ 0.3). On independent datasets, performance collapsed and became worse than a baseline model that always predicted the training set mean (*R*^2^ < 0). We found that these failures were not due to model complexity or measurement noise, but rather to a fundamental problem: covariate shift. A lack of shared taxa, differences in sample preservation, and even differences in taxonomic classifier versions between training and test data derailed models which relied on fixed input formats and fragile assumptions.

Covariate shift is a widespread yet underappreciated challenge for the many machine learning models that rely on microbiome taxonomic features (e.g., Lo and Marculescu ^36^, Metwally et al. ^37^, Sharma and Xu ^38^, Armour et al. ^39^, Kubinski et al. ^40^, Jurvansuu et al. ^41^, Chang et al. ^42^, Kosciolek et al. ^43^, Chen et al. ^44^, Ananthakrishnan et al. ^45^). Beyond the lack of shared taxa between studies, taxonomic features derived from sequencing data are highly sensitive to reference databases, classifier versions, and pipeline-specific choices. As these tools evolve, the set of taxa detected can shift dramatically, making it difficult to ensure that model inputs remain compatible over time. Models that expect fixed feature sets can fail silently when confronted with mismatched or incomplete inputs—for example, by replacing missing taxa with zeros or ignoring new taxa entirely. These behaviors can cause predictions to collapse or become erratic, especially when the model lacks built-in safeguards or input validation. Without rigorous benchmarking and transparency, even well-intentioned tools risk producing misleading or irreproducible results when deployed outside the context they were trained on.

One reason these limitations may have been overlooked is that model performance was evaluated primarily using sample correlation. While sample correlation measures the strength of association, it can be near one when the slope of the relationship is far from one–meaning that models that predict only within a narrow range of values can still appear to perform well. In contrast, the coefficient of determination (*R*^2^) quantifies the proportion of variance explained by the model, penalizing both biased and low-variance predictions. This distinction is critical in microbiome studies, where microbial load estimates are used in downstream analyses that are sensitive to such biases^9^. We therefore recommend that future work in this area avoid reliance on sample correlation and instead adopt more appropriate metrics such as *R*^2^.

Our results provide the largest systematic study of scale to date and add to the growing literature advocating for explicit modeling of scale uncertainty in microbiome analyses. Like the ML prediction models of Nishijima et al. ^15^, normalization methods assume that microbial load can be imputed without error from sequence count data^9^. Ignoring error makes these methods fragile, leading to biased estimates and misleading scientific conclusions. In contrast, Bayesian PIMs explicitly represent uncertainty in scale, yielding more robust and reproducible inference^46^. Across 30 benchmark datasets, we find that Bayesian PIMs substantially improved false discovery control without compromising power.

In this study, we deliberately used a simple Bayesian PIM based on a weakly informative scale model reflecting broad biological plausibility, chosen to be valid across diverse datasets. The real strength of the framework, however, emerges when more informative scale priors are specified. Prior work has shown that incorporating expected shifts in microbial load—such as decreases after antibiotic exposure or increases during ecological colonization—can substantially increase statistical power while maintaining nominal false discovery rates^7,9^. Scale models can also be anchored by external measurements like flow cytometry or qPCR, while still accounting for their uncertainty^7,9^. Such context-aware models let researchers inject domain knowledge without overcommitting to unverifiable assumptions, offering a flexible and interpretable path forward for robust microbiome analysis.

Importantly, these conclusions hold even if future machine learning models achieve higher accuracy in predicting microbial load. Such predictions could serve as informative priors or inputs within Bayesian scale models, where their uncertainty can be formally quantified and propagated into downstream analyses. The goal is not to dismiss machine learning, but to integrate it into frameworks that respect the fundamental identifiability limits of sequencing data–ensuring that advances in prediction strengthen, rather than undermine, robust inference.

To support the development and evaluation of these methods, we curated *mutt*, the largest public repository of microbiome datasets with paired sequencing and microbial load measurements. By providing processed and reprocessed data, standardized metadata, and open tools, *mutt* lowers the barrier to reproducibility research and enables meaningful benchmarking across diverse settings. We invite the community to build on this resource, contribute additional datasets, and adopt reproducible practices in model development.

Finally, we emphasize a broader lesson for microbiome modeling: statistical identifiability is not a nuisance to be ignored, but a reality to be respected. Modeling scale is hard not because the models are underpowered, but because the data do not contain the necessary information. Rather than chasing ever more complex models trained on fragile inputs, we advocate for methods that model what can–and cannot–be known. In microbiome data analysis, uncertainty is not the enemy. It’s the path to robustness.

### Limitations of the study

While we curated and screened over 120 studies, many human gut microbiome datasets with paired microbial load measurements remain inaccessible due to data-sharing restrictions or lack of deposited raw data. Additionally, while we attempted to evaluate other ML-based scale prediction tools (e.g., Wirbel et al. ^47^), software and data limitations prevented execution or proper comparison. These models may offer improvements, especially for specific contexts like stool samples or extraction protocols, but rigorous cross-study validation is still needed.

## RESOURCE AVAILABILITY

### Lead contact

Requests for further information and resources should be directed to and will be fulfilled by the lead contact, Justin Silverman.

### Materials availability

This study did not generate new unique reagents.

### Data and code availability

- All data used in this manuscript have been previously published and are publicly available. A list of project accessions are deposited in Supplemental Table 1. All data and issue tracking for the data analyzed in this study are openly available for download at https://github.com/Silverman-Lab/mutt using the publicly available R package *mutt*. The README available details a step-by-step vignette on how to contribute and utilize each feature of the data repository to ensure reproducibility.
- All original analysis code is available on https://github.com/maxwellkonnaris/mlscale and is publicly available as of the date of publication.
- Any additional information required to reanalyze the data reported in this paper is available from the lead contact upon request.

## ACKNOWLEDGMENTS

We thank Dr. Rachel Silverman and Dr. Travis Gibson for their manuscript comments. Also, we thank Won Gu and Tinghua Chen for contacting authors to acquire publicly available data. MK was supported in part by the National Institute of Diabetes and Digestive and Kidney Diseases (NIH 5T32DK120509-05). JDS was supported in part by the National Institute of General Medical Sciences (1R01GM148972-01).

## AUTHOR CONTRIBUTIONS

Conceptualization, MAK and JDS; methodology, MAK and JDS; investigation, MAK and JDS; data acquisition, MAK and MS; data processing, MAK and MS; data curation, MAK; software, MAK; formal analysis, MAK; validation, MAK and JDS; visualization, MAK; writing–original draft, MAK and JDS; writing–review & editing, MAK, MS, NL, and JDS; funding acquisition, MAK and JDS; resources, JDS; project administration, JDS; supervision, JDS.

## SUPPLEMENTAL INFORMATION INDEX

Document S1. Figures S1–S2.

Table S1: Studies surveyed and collected for *mutt*, studies included in analysis, studies included in differential abundance, related to Figure 1, 2, 3.

Table S2: Performance metrics, study characteristics, and author presence for each dataset in the analysis, related to Figure 1, 2, and 3.

Table S3: Differential abundance results, related to Figure 3.

## STAR METHODS

### Key resources table

#### Experimental model and study participant details

##### Database Construction

PubMed and Google Scholar were queried for studies published between 2009 and 2025 that reported absolute abundance data. Searches used combinations of the keywords *microbiome, absolute abundance, quantitative microbiome profiling*, and *total microbial load*, together with measurement modalities (*flow cytometry, ddPCR, qPCR, spike-in DNA*) and sequencing strategies (*amplicon — 16S, ITS, 18S rRNA, shotgun metagenomics*). Studies without paired microbial profile and load estimates were excluded, leaving approximately 120 studies. Each first or corresponding author, as well as any designated point of contact, was contacted at least twice, with follow-ups spaced at approximately 3-week intervals.

Where raw sequencing data could be obtained count tables were reprocessed to be consistent with the Nishijima et al. ^15^ study. Amplicon datasets were denoised with DADA2^20^ and taxonomically classified using the RDP Classifier (v16)^48^. Shotgun metagenomic datasets were profiled with MetaPhlAn4^19^, using single-stranded input when applicable. Resulting merged count matrices for each dataset were subsequently converted to proportions. A continuously updated index of studies reporting absolute microbial abundances is maintained in the *mutt* repository, and the full list of studies analyzed here is provided in Supplemental Table 1.

##### Study and Sample Inclusion

In addition to the four studies^6,15,17,18^ included in Nishijima et al. ^15^, 31 External studies were selected from the *mutt* repository for inclusion in this analysis. Each study contained either 16S rRNA amplicon or shotgun metagenomic sequencing data paired with flow cytometry, ddPCR, or qPCR measurements of total microbial load^3,23–29,49–71^.

Studies were excluded if they relied solely on spike-in standards, colony-forming unit (CFU) counts, mock community controls, or ITS/18S rRNA sequencing. Studies still undergoing preprocessing or validation at the time of writing were also excluded. When both original and reprocessed sequencing data were available for a given study, the version yielding the highest predictive model performance was selected (i.e., a best-case scenario). In cases with multiple external load measurements of the same modality (e.g., average cells per microliter versus total cells per five minutes by flow cytometry), the modality that maximized model performance was retained.

### Quantification and statistical analysis

#### Application of the *Microbial Load Predictor*

The GALAXY, MetaCardis, and Vandeputte (2021) pretrained models from Nishijima et al. ^15^ were accessed via the *Microbial Load Predictor* library downloaded on June 18, 2025 from (https://github.com/grp-bork/microbial_load_predictor/). We used the 16S rRNA models (Vandeputte [2021]) trained on RDPv16 classified data. We used the metagenomic models (GALAXY and MetaCardis) trained on data processed using MetaPhlAn4(metaphlan4.mpa_vJun23_CHOCOPhlAnSGB_202403) for the analysis of all metagenomics datasets less the GALAXY and MetaCardis datasets which were analyzed using the mOTUv2.5 processed data to reproduce the analysis of Nishijima et al. ^15^. Prior analyses have shown only minimal differences in model performance between these two processing methods^72^. Relative abundance tables were processed following the methods of Nishijima et al. ^15^.

#### Model Evaluation

External microbial load measurements obtained via flow cytometry, ddPCR, or qPCR were treated as the ground truth for evaluating model predictions. As the originally published *Microbial Load Predictor* reported outputs in log_10_ scale, both predicted and measured microbial loads were transformed accordingly. Let *y* = (*y*_1_, …, *y*_*n*_) denote the vector of measured microbial load, where each *y*_*i*_ corresponds to the external measurement (e.g., via flow cytometry, ddPCR, or qPCR) for sample *i* and let 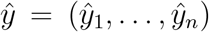 denote the vector of predicted microbial load, where each 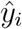 is the output of the *Microbial Load Predictor* for sample *i*. Performance was then evaluated using the correlation between measured and predicted microbial load on a log scale.

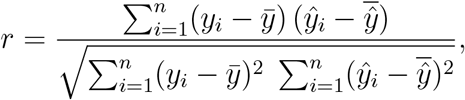

where 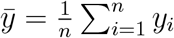 and 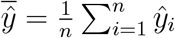.

Correlation, *r*, can be near one even when the slope of the relationship between two variables is far from one–e.g., when a predictive model predicts in a restricted range. By contrast, the coefficient of determination (*R*^2^) does not have this problem. We adopt the variance-partitioning definition: 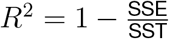 where SSE is the sum of squared errors and SST is the sum of squares of the true values. *R*^2^ measures the proportion of variance in the true values explained by the predictions themselves. Because absolute microbial load estimates can differ systematically in scale across measurement modalities, both predicted and true loads were mean-centered before computing *R*^2^ to prevent global offsets from artificially deflating performance estimates:

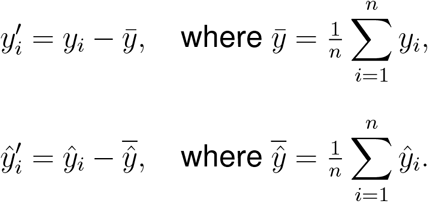

This adjustment yields a conservative, modality-agnostic measure of explained variance:

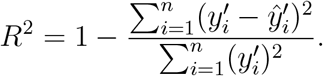

Here, *R*^2^ = 1 denotes perfect prediction, *R*^2^ = 0 indicates no improvement over predicting the mean, and negative values imply worse performance than a baseline predictor that always predicts the training set mean.

#### Visualizing covariate shift using dimension reduction

For each sample, the relative abundance profile was filtered to retain only the features used by the *Microbial Load Predictor* 16S rRNA model (the Vandeputte2021 model). Features absent from a given dataset were added and imputed with zeros, following Nishijima et al. ^15^ ‘s preprocessing procedure. All study datasets were merged, and a t-SNE embedding was computed using the `Rtsne` package to obtain a low-dimensional representation of feature space. To visualize the distribution of training samples, we overlaid a 99% confidence ellipse computed using the `stat ellipse` function in `ggplot2`. This ellipse is derived from the sample mean and covariance of the Vandeputte (2021) training data in the t-SNE space and represents the region expected to contain approximately 99% of training samples.

#### Benchmarking differential abundance analysis

The Scale Reliant Inference (SRI) framework^9^ expresses the absolute abundance of taxon *d* in sample *n* as the product of its composition (proportional abundance; 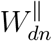) and scale (total microbial load; 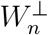:

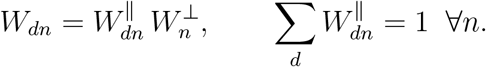

Here, 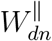 denotes the proportional abundance of taxon *d* in sample *n* and 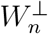 the total microbial load of sample *n*.

Under this factorization, the log_2_ fold change (LFC) in absolute abundance between two conditions decomposes into compositional and scale components:

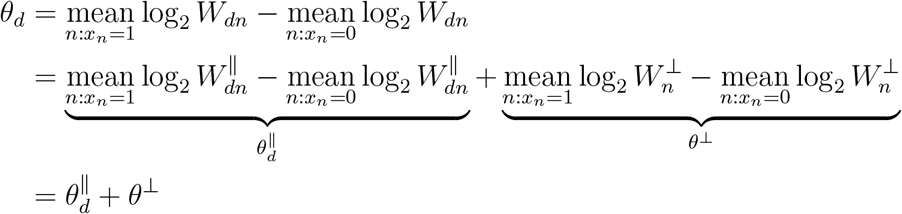

where *x*_*n*_ is a binary condition indicator, 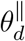 captures changes in relative abundance between conditions, and *θ*^*⊥*^ captures changes in total microbial load. Detecting differential abundance therefore requires estimating both terms. For example, if a taxon’s relative abundance doubles 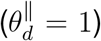 while total microbial load halves (*θ*^*⊥*^ = −1), the net change is *θ*_*d*_ = 0. Ignoring scale in this case could lead to the mistaken conclusion that the taxon was decreasing, simply because its relative abundance declined.

One of the central tools in SRI is the Bayesian PIM, specified in a factorized form called a *Scale Simulation Random Variable (SSRV)*. An SSRV has modular components: a measurement model *p*(*W* ^∥^ | *Y*), a scale model *p*(*W* ^*⊥*^), and an estimand (here *θ*_*d*_). Together, these define a Bayesian PIM (see Nixon et al. ^9^ for details). In this article, we consider the following scale models:

- **External load measurements**. If microbial load is directly measured (e.g., by flow cytometry, ddPCR, or qPCR), then we follow Nixon et al. ^7^ and use

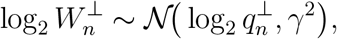

where 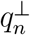 is the measurement for sample *n* and *γ*^2^ reflects potential measurement error variance.

- **Normalization as a scale model**. In the absence of external data, normalization implicitly defines a scale model^9^. In this article we use Total Sum Scaling (TSS) which is a special case of the following scale model

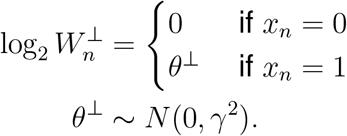

TSS normalization is obtained in the limit as *γ* → 0^8^. Intuitively, this corresponds to assuming no change in microbial load between conditions (*θ*^*⊥*^ = 0); allowing *γ* > 0 relaxes this assumption by modeling uncertainty in potential shifts.

- **Machine learning predictions**. The Nishijima et al. ^15^ approach estimates relative abundances from sequence count data and then scales those relative abundance by model predicted microbial loads. Within the SSRV framework, this is analogous to the following scale model

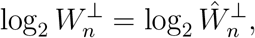

where 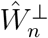 denotes the model prediction for the *n*-th sample. That is, this approach does not consider potential error in the predicted microbial load.

We implemented these alternatives within ALDEx3, a rewrite of ALDEx2 with improved computational efficiency and a streamlined model interface^46^. As in ALDEx2, Monte Carlo replicates of the count table are simulated from a Dirichlet–multinomial to capture sampling variability (the measurement model, *p*(*W* ^∥^ | *Y*)). Each replicate is rescaled using a sample from the chosen scale model (e.g., TSS, external measurements, or ML predictions), and LFCs are then estimated on absolute abundances. Hypothesis testing uses Wilcoxon rank-sum tests, with results averaged across replicates to propagate uncertainty from both composition and scale^7^.

We benchmarked ANCOM-BC2^32^, DESeq2^30^, Limma-Voom^31^, edgeR^33^, metagenomeSeq^34^, LinDA^35^, ALDEx3 under TSS normalization with and without added scale uncertainty (*γ* = 0, 1 in log_2_ units)^7^, and ALDEx3 with ML-predicted scale^15^. Analyses included *N* = 24 amplicon and *N* = 6 shotgun metagenomic datasets with paired counts and metadata. Ground truth was defined from external microbial load measurements, log_2_-transformed and treated as the mean of a normal distribution with standard deviation 0.5 to account for measurement uncertainty as suggested by Nixon et al. ^7^. Fold-change direction and adjusted *p*-values (Benjamini–Hochberg^73^) were compared against this ground truth to calculate true and false positives/negatives. From these, we derived Positive Predictive Value (PPV), Negative Predictive Value (NPV), and False Discovery Rate (FDR) as benchmarks of performance.

