## Supplementary material for "Uncertainty Modeling Outperforms Machine Learning for Microbiome Data Analysis": DocumentS1

### Table of Contents

S1 Sample correlation can misrepresent predictive performance. . . . . . . . . . . . . . 2
S2 Prediction variance collapse manifests as high mode frequency within studies. . . 3

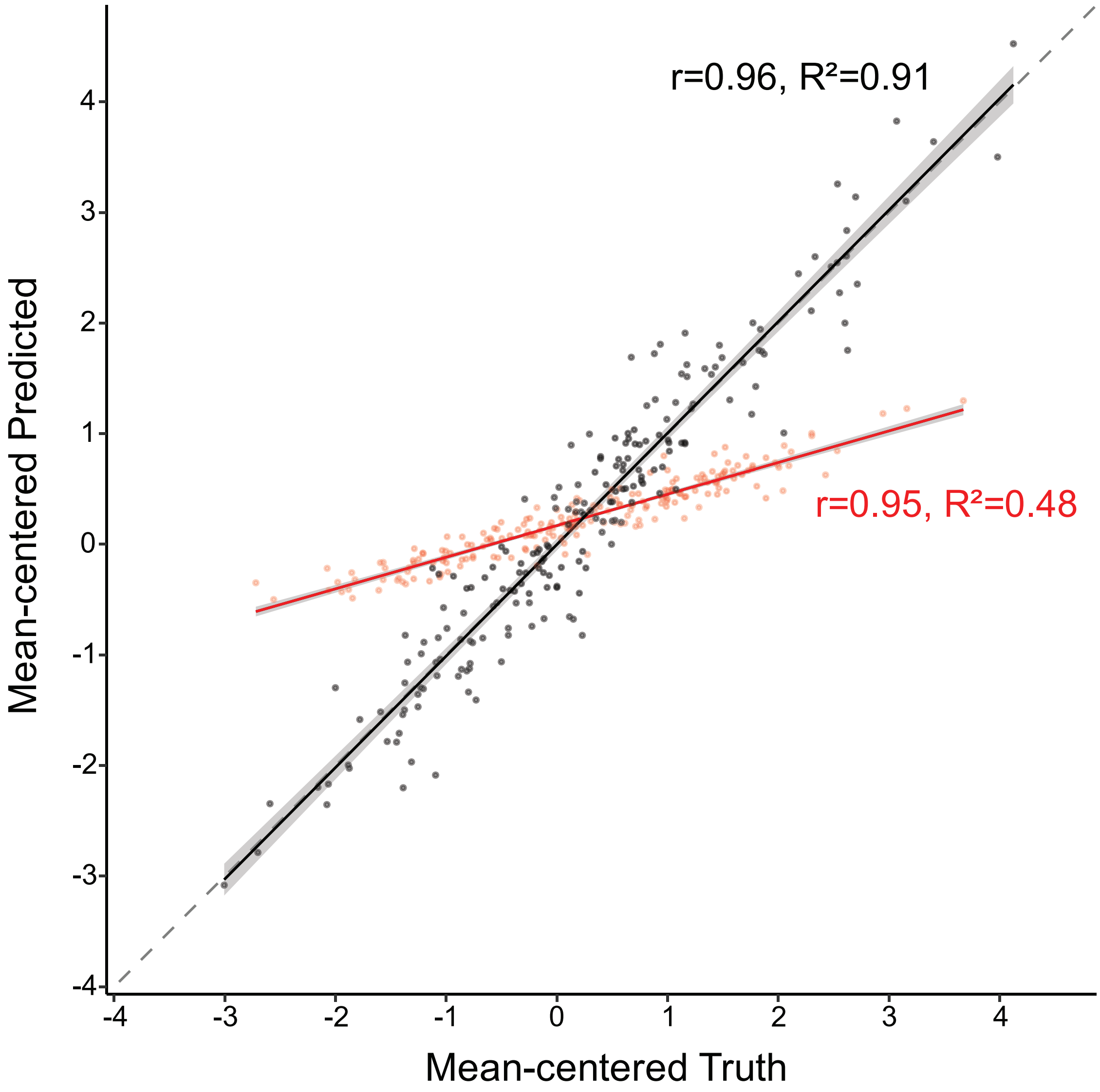


**Figure S1. Sample correlation can misrepresent predictive performance.** Two predicted models were simulated, one (red) that significantly underestimates the range of the target and one (black) that provides more accurate predictions that match the range of the target. Sample correlation $r$ can be near one even when the predictions have a much narrower range than the true target values. In contrast, the coefficient of determination $R^2$ is only near one when there is a near one-to-one correspondence between predictions and the truth.

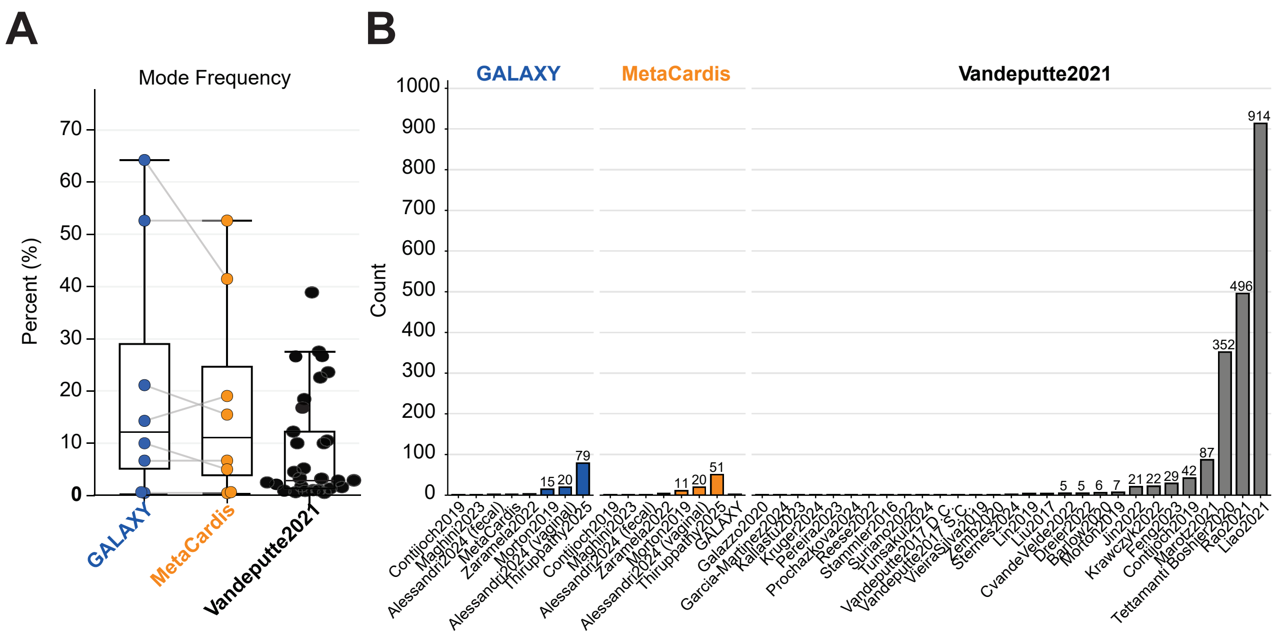


**Figure S2. Prediction variance collapse manifests as high mode frequency within studies. (A)** Mode frequency—the proportion of samples assigned the single most common predicted value—is shown for each external study using the GALAXY-trained model (blue), the MetaCardis-trained model (orange), and the Vandeputte (2021) model (black). Lines connect results from the same study across the two metagenomic models to illustrate variability. High mode frequency indicates that a large fraction of samples received essentially identical predictions, reflecting a collapse of variance and failure to capture biological differences. **(B)** The corresponding total number of samples assigned to the mode is shown for each study. Annotations are provided only for cases where the mode comprised more than four samples; in some studies, hundreds of samples shared an identical predicted value. Together, these panels show that the Nishijima *et al.* models often default to a single prediction across most samples within a study, rather than generating meaningful sample-level variation.
